# Fortune may favor the flexible: Environment-dependent behavioral shifts in invasive coquí frogs

**DOI:** 10.1101/2023.10.30.564823

**Authors:** Katharina M Soto, Devin Edmonds, Andrea L Colton, Faith O Hardin, Eva K Fischer

## Abstract

Biological invasions are a major driver of global biodiversity loss, impacting endemic species, ecosystems, and economies. While the influence of life history traits on invasive success is well-established, the role of behavior in the invasive potential of animals is less studied. The common coquí frog, *Eleutherodactylus coqui*, is a highly successful invader in Hawaii. We build on previous research characterizing changes in physiology and morphology to explore behavioral variation across the invasive range of coquí in Hawaii. We hypothesized that behavioral traits contributing to invasion expansion would be reflected in behavioral differences at the invasive center versus invasive edge. To investigate whether differences in the field represent local adaptation or behavioral plasticity, we additionally evaluated behavior following acclimation to a shared laboratory environment. While we identified only subtle behavioral variation among populations in the field, we found that individuals from all populations became less bold, active, and exploratory in the laboratory, converging on a similar behavioral phenotype. Alongside previous work, our results suggest that coquí adjust their behavior to local environmental conditions across their invasive range, and that behavioral flexibility may contribute to invasive success.

## Introduction

Biological invasions are a major cause of global biodiversity loss, impacting endemic species, ecosystems, and economies (Kumschick et al. 2012; Lennox et al. 2015). While the impacts of invasive species have been thoroughly examined, we are still in the process of understanding the attributes that contribute to invasive species’ success. Research examining invasive animals has demonstrated the importance of life history traits, such as high reproductive rates (Alcaraz et al. 2005), rapid population growth, dispersal capabilities, and a generalist diet (Sakai et al. 2001; Alex Perkins et al. 2013). The role of behavior remains less studied, but has been implicated in the spread of some invasive species (Sol and Lefebvre 2000; Weis 2010; Hudina and Hock 2012; Damas-Moreira et al. 2019). For instance, Candler & Bernal (2015) found that introduced cane toads (*Rhinella marina*) displayed reduced neophobia and increased exploration of novel prey and objects compared to toads from native populations (Candler and Bernal 2015). Similarly, Damas-Moreira et al. (2019) found that invasive wall lizards (*Podarcis sicula*) were more bold, exploratory, and neophilic compared to non-invasive wall lizards (*Podarcis virescens*) (Damas-Moreira et al. 2019). These studies highlight how certain behavioral traits and/or collections of behaviors (“behavioral syndromes”) can drive invasive success by facilitating range expansion and the exploitation of new resources. Furthermore, behavioral plasticity may be particularly crucial during the initial stages of invasion when adaptation can be slow due to low genetic diversity resulting from founder effects and bottlenecks during colonization (Tsutsui et al. 2000; Peacock et al. 2009; Wright et al. 2010). Yet to succeed, invaders must adjust rapidly to novel environments, prey, predators, and competitors (Chapple et al. 2012), and behavioral plasticity may help bridge this gap.

Behavioral and physiological changes may be closely linked to invasion potential as behavior (behavioral plasticity) may help overcome novel physiological challenges, and physiological adjustments may in turn facilitate coordination of favorable behavior. Indeed, colonization of novel environments often presents physiological challenges, and behavior may be an important coping mechanism. High metabolic rates, often associated with a ’fast pace of life’—marked by rapid growth, early reproduction, and a generalist diet—are traits already recognized as key contributors to invasive potential (Lagos et al. 2017). Additionally, changes in metabolic rates have been correlated with changes in boldness, exploration, and activity (Biro and Stamps 2010; Myles-Gonzalez et al. 2015; Baškiera and Gvoždík 2022), behaviors that have been identified as relatively greater in invasive species as compared to native organisms (Candler and Bernal 2015; Damas-Moreira et al. 2019). These relationships may be particularly important in ectotherms that directly regulate body temperature through their behavior; however, connections between behavior and physiology during biological invasions have only been examined in a few systems (Myles-Gonzalez et al. 2015; Mathot et al. 2019; Mowery et al. 2021).

The common coquí (*Eleutherodactylus coqui*) is an excellent model for investigating mechanisms that contribute to invasion success. These small, nocturnal frogs are endemic to Puerto Rico, where they are an integral part of the island’s ecosystem and culture (reviewed in Westrick et al. 2022). However, their introduction to other regions has garnered them a reputation as one of the worst invasive species in the world (Lowe et al. 2000). Coquí have successfully colonized Hawaii, Guam, and various Pacific and Caribbean islands (Kraus and Campbell 2002; Beard and Pitt 2005), and have been introduced to Costa Rica (Barrantes-Madrigal et al. 2019). Their invasion of Hawaii extends across the archipelago, including the islands of Hawaii, Maui, Oahu, and Kauai (Beard et al. 2009), though the Kauai population is believed to have been exterminated (Beard and Pitt 2012). Their loud and incessant calls have made them an unmistakable presence in the Hawaiian nightscape, impacting property values, tourism, and the ornamental plant trade, and prompting campaigns to prevent their spread (Beard and Pitt 2005).

Since their introduction to Hawaii in the 1980s, coquí populations have exploded to densities three-fold those in their native range, reaching up to 91,000 frogs per hectare (Beard et al. 2009). Unlike most frogs, coquí are direct developing, meaning they complete metamorphosis within the egg, skipping a free-living tadpole stage. This life history allows for rapid reproduction and independence from standing freshwater (reviewed in Westrick et al. 2022). This reproductive strategy, combined with a voracious, generalist diet, has fostered their rapid population growth.

On the Big Island of Hawaii where populations are largest, coquí are continuing to expand, including to higher elevations (Marchetti et al. 2023; Marchetti et al. 2024). Generally, this combination of factors has led to a tradeoff between escaping high competition at low elevations and coping with colder and more variable environmental conditions further uphill. Previous studies have documented the invasive spread of coquí in Hawaii (Marchetti et al. 2023; Marchetti et al. 2024), characterized the impacts of coquí on local plants, animals, and economies (Kraus and Campbell 2002; Beard and Pitt 2005; Beard et al. 2008; Smith et al. 2018), and explored physiological changes associated with the colonization of distinct habitats (Beard et al. 2008; Rollins-Smith et al. 2015; Haggerty 2016; O’Neill et al. 2018; Marchetti et al. 2023; Marchetti et al. 2024). Behavior remains relatively understudied, aside from Marchetti and Beard (2021) who demonstrated that Hawaiian coquí maintain avoidance responses to native Puerto Rican predators ∼20 generations after invasion. Like other invasive species, Hawaiian coquí exhibit reduced genetic diversity as compared to their native counterparts due to founder effects (Tsutsui et al. 2000; Peacock et al. 2009). Nonetheless, their populations have thrived, establishing a presence in diverse habitats and ecosystems across the island. This versatility in the face of low genetic diversity suggests a role for plasticity in their invasive potential (Wright et al. 2010). Indeed, the above studies document a combination of population differences and plasticity in various traits.

Our goal was to investigate how behavior might contribute to the invasive expansion and success of *E. coqui* in Hawaii. We adopted a stepwise approach, involving field and laboratory experiments. First, we characterized activity, exploration, and boldness across elevational and density gradients in Hawaii to understand to what extent behavior varies across environmental conditions at the invasive center verses edge (Chuang and Peterson 2016). We considered high-density, low-elevation sites representative of the invasive center, and low-density sites at both high and low elevations as invasive edges. We tested the alternative predictions that (1) more active and bold coquí would be found at invasive edges as these individuals would be more likely to push range boundaries, or (2) more bold, active coquí would be found at the invasive center as these individuals would be better equipped to deal with the competitive challenges of high densities. By targeting high- and low-density sites at different elevations, we additionally asked whether the effects of density were modulated by more challenging environmental conditions.

Second, we asked whether behavioral traits changed when we introduced coquí to a novel, laboratory environment, specifically measuring behavior in the same individuals in the field and in the laboratory. This allowed us to distinguish environmentally-mediated behavioral plasticity from stable behavioral differences that would indicate developmental influences or local adaptation to different densities and/or elevations. We also measured resting metabolic rate in the laboratory to test for an association between metabolic rate and behavior, thereby exploring a potential mechanism by which behavioral changes are coordinated and linked with physiological adaptation to high elevations. Taken together, our findings contribute a behavioral perspective to complementary studies characterizing life history and metabolic traits in Hawaiian coquí, adding an additional dimension to our understanding of this small frog’s invasive success.

## Methods

All frog husbandry and experimental methods were approved by the University of Illinois Urbana-Champaign Animal Care and Use Committee (Protocol # 20147). Frog collection and export were approved by the State of Hawaii Department of Land and Natural Resources (Permit # H0422-04; EX-22-11). Given the invasive nature of coquí in Hawaii, our collection posed no conservation concerns.

### Field Collection

We collected 60 male coquí on the Big Island of Hawaii. We collected frogs from four site types: high-density/low-elevation (N=16), low-density/low-elevation (N=14), high-density/high-elevation (N=15), and low-density/high-elevation (N=15). Collection sites were at Lava Tree State Monument Park (HL, 195m, [19°28’58“N 154°54’09”W]), ’Ōla‘A Forest Reserve (LH, 870m, [19°27’10“N 155°11’14”W]), Watershed Forest Reserve (LH, 860m, [19°25’02“N 155°09’45”W]), Upper (HH, 850m, [19°34’28“N 155°11’27”W]) and Lower (HL, 250m, [19°37’46“N 155°05’37”W]) Stainback Highway, Homestead Road (LL, 200m, [19°50’33“N 155°06’50”W]), and Hilo Watershed Reserve (HH, 720m, [19°50’36“N 155°11’08”W]).

We defined low-elevation as <300M and high-elevation as >700m. We classified population density based on recent surveys (Marchetti et al. 2024), and used call surveys during collection as a rough confirmation of density differences, considering higher sound pressure is associated higher male coquí density (Benevides et al. 2019). We performed acoustic recordings at each collection site and our field station using a shotgun microphone (Sennheiser ME 66/K6) and a handheld recorder (Zoom H1 Handy Recorder). Intensity analysis of recorded calls was conducted using Audacity v. 3.0.2 (Audacity Team 2021). **L**ow-density sites had an average frequency of 500 Hz at 40.7 dB and high-density sites had an average frequency of 2,639 Hz at 55.6 dB, lending support to our initial classifications based on Marchetti et al. 2024.

Frogs were hand-captured between 19:00 and 00:00 hours and individually placed in 500 mL plastic cups with ventilation holes and damp paper towels for moisture and humidity. Frogs were transported from each collection site to our field station in Leilani Estates, Hawaii County, HI, and held for 24 hours before behavioral testing the following night. We conducted all field behavioral assays in this manner to minimize the acute effects of temperature and humidity differences between sites on behavior, make the timing of trials uniform, allow frogs to recover from capture stress, and due to limitations in transporting our behavioral arena and recording equipment.

### Field Behavioral Assays

We conducted field behavioral trials between May and June 2022, from 19:00 to 03:00 hours. Average temperature was 22.9 °C (range: 19–25) and humidity was 76.7% (range: 70–85). We used two behavioral arenas constructed from ½” PVC pipes and white mesh fabric, with dimensions measuring 83 x 58 x 48 cm. The arenas were illuminated by red light lamps and infrared illuminators (Univivi IR Illuminator), and we recorded behavior using infrared camcorders (Sony DCR-SR85). Arenas were placed ∼8 meters apart on opposite sides of the field station to prevent trial interference. We constructed emergence chambers using a ½” PVC pipe held vertical by a plastic cup base and a retractable cardboard cover.

To examine behavior, we used two tests: (1) an emergence test to quantify latency to emerge from a shelter (a commonly used metric of boldness, see Magnhagen et al. 2014; Myles-Gonzalez et al. 2015; Yuen et al. 2017), and (2) an open field test to quantify activity and exploration. For the emergence test, each frog was placed in the emergence chamber and given a 3-minute acclimation period. After the acclimation period, the cover of the chamber was removed from outside the arena. Each frog was given 10 minutes to emerge from the chamber.

Individuals that did not emerge after 10 minutes were gently removed from the PVC pipe without direct handling of the frog. To ensure this experience did not bias subsequent behavior, we compared activity and exploration between frogs that emerged on their own versus those that were removed and found no differences (exploration: F_1,52_= 0.04, p= 0.838; activity: F_1,52_= 0.06, p= 0.801). Latency to emerge was recorded as the moment when the majority of the frog’s body had exited the PVC pipe. Immediately following the emergence test, we performed an open field test for 15 minutes. During the open field test, we quantified exploration as the proportion of areas visited, and activity as the number of times a frog crossed lines between areas superimposed during analysis (see below). We expected a positive correlation between activity and exploration; however, these two variables are not perfectly correlated and can vary between individuals or populations, as frogs that move the same amount can do so over smaller or larger areas (e.g., 10 line crossings back and forth between 4 areas versus 10 line crossings across 10 areas).

### Laboratory Transfer and Husbandry

Within four days of capture, all frogs were shipped to the University of Illinois Urbana-Champaign. For transport, each frog was placed in a 500 ml plastic cup with a moist paper towel and dead leaves. Cups were packed into Styrofoam boxes with insulation and padding material and shipped via climate-controlled UPS overnight. All frogs arrived alive and in good body condition. Upon arrival, frogs were individually housed in tanks measuring 30 x 20 x 15 cm, with ∼ 2-3 cm of soil and sphagnum moss as a substrate. Additionally, each frog was provided a ∼20cm long PVC pipe as a shelter. We maintained humidity >70% and temperature 69–74 °F on a 12L:12D light cycle with a shifted dark period (lights off at noon and on at midnight). We fed frogs live brown crickets (*Acheta domesticus*) gut loaded with vitamins three times weekly. Four individuals died over the course of the study due to escape and subsequent desiccation.

### Laboratory Behavioral Assays

We conducted laboratory trials (N=56) between September and November 2022 after frogs were confirmed healthy and had acclimated to the laboratory for 14–18 weeks. All trials were conducted between 13:00 and 17:00 hours, one to four hours into the frogs’ wake cycle after lights went off. To replicate the conditions of field capture and acclimation, we held individuals in 500 mL plastic cups lined with moist paper towel for 24 hours prior to laboratory behavioral trials. Trials were conducted in an unoccupied animal room with average temperature 21.7 °C (range: 18–24) and average humidity 74.8% (range: 50–98). We used the same arena as in the field assays; however, we used curtains to dampen sound reflections off the walls and maintain humidity in the arena. Behavioral assays in the laboratory closely followed the field protocol, consisting of a 13-minute emergence test (3 minutes acclimation and 10 minutes to emerge) immediately followed by a 15-minute open field test. Ambient sound recordings collected during behavioral trials in the field were used to replicate natural background noise, including the presence of other coquí.

### Physiological Assays

We conducted resting metabolic rate (RMR) assays (N=55) in the laboratory between February and March of 2023, 12 weeks after laboratory behavioral assays. Given that resting metabolic rate (RMR) can be influenced by numerous factors, including diet, temperature, age, sex, and environmental conditions (particularly in ectotherms), prior studies have highlighted that a common garden approach is the preferred method for controlling these variables (Biro and Stamps 2010; Burton et al. 2011).

We used an intermittent flow-through respirometry system to measure RMR. We placed frogs in 127 mL chambers made from ½” diameter PVC plastic pipe and PVC end caps. We configured a flow-through system with an open circuit and incurrent flow measurement using a PP2 Dual Pump, FB8 Flow Measurement System, RH-300 Water Vapor Analyzer, Fox Box, and UL2 Universal Interface (all from Sable Systems International). The incurrent airstream was scrubbed of CO_2_ and water vapor. After allowing frogs to acclimate in the chamber for 20 minutes, we sampled CO_2_ production and O_2_ consumption for each individual every 5 seconds over a 40-minute window. We switched from a chamber containing a frog to a control chamber between individuals to establish a baseline measurement of ambient CO_2_ and O_2_, and to account for measurement drift. Measurements were recorded following a 2-day fast to eliminate the energy expenditure associated with digestion (Naya et al. 2009). To calculate RMR, we first matched each frog to the control measurement taken directly after the trial. For three individuals, we substituted controls from the following trial on the same day due to problems with their paired control. We deleted 5 minutes from the start of each trial to remove any error associated with switching tubing between the control and frog. We also deleted 7.3 minutes (determined from visually inspecting plots) from the end of each trial to remove the lag between sequential CO_2_, O_2_, and H_2_O sensors in our system. After trimming data, we calculated the fractional concentration of CO_2_ and O_2_, and H_2_O adjusted for barometric pressure.

To calculate RMR, we first calculated the rate of O_2_ consumption (*VO*_2_) and CO_2_ emission (*VCO*_2_) using Lighton (2008) equations 10.1:

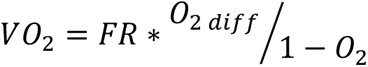

and 10.8:

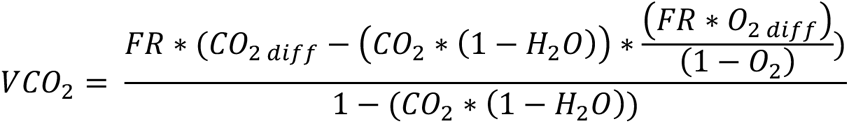

where *FR* = flow rate, *O*_2*diff*_ is the average O_2_ during the control minus the O_2_ produced by a frog, *CO*_2*diff*_ is the CO_2_ produced by a frog minus the average CO_2_ during the control, and *CO*_2_, *O*_2_, and *H*_2_*O* are the average values during control. We then calculated RMR as the lowest 3-minute average *VO*_2_ and *VCO*_2_ during each trial. Next, we calculated the respiratory quotient (RQ) by dividing the lowest 3-minute average *VCO*_2_ by the lowest 3-minute average *VO*_2_. Finally, we calculated mass corrected RMR by dividing each frog’s RMR by its mass (g), hereafter referred to as RMR. Data manipulation was performed using the packages ‘zoo’ and ‘tidyverse’ (Wickham et al. 2019; Zeileis et al. 2023) in R.

### Statistical Analysis

We used Behavioral Observation Research Interactive Software (BORIS) version 8.13 (Friard and Gamba 2016) to code exploration and activity from videos. We analyzed data in R version 4.2.4 (R Core Team 2022) in Rstudio v.4.2.2 (RStudio Team 2021). We analyzed response variables (i.e., behavior and RMR) using raw data for those variables with a normal distribution. We applied a square root transformation to activity because it was not normally distributed. We lost field video recordings for five frogs from low-density/low-elevation (LL) and eight frogs from high-density/low-elevation sites (HL) due to a hard drive failure. The analysis of field behavior (i.e., exploration and activity) is based on the remaining dataset (N=49). Field emergence data was recorded live during trials, and all available data (N=60) were analyzed. We used power analysis with the ‘pwr’ package (Champley 2020) in R to assess the consequences of this data loss. Even with reduced sample sizes, we maintained high power (94%) to detect large effects and moderate power (64%) to detect medium effects. We used the R packaged ‘effectsize’ (Ben-Shachar et al. 2020) to estimate eta-squared (η^2^) effect sizes.

To examine behavioral differences in the field, we used the ’lm’ function from the ’stats’ package to construct linear models that included behavior (activity or exploration) as the response variable, and density, elevation, and their interaction as predictor variables. We used similar models to test for behavioral and RMR differences in the laboratory. For linear models, statistical significance was called at p ≤ 0.05 and a trend toward statistical significance at p ≤ 0.10. We interpreted effect sizes: η^2^ > 0.01 as a small effect, η^2^ > 0.06 as a medium effect, and η^2^ > 0.14 as a large effect,

For emergence, not all frogs performed the behavior during the trial period, making a direct test of latency to emerge problematic. To address this issue, we ran Cox Proportional Hazard Models using the R packages ‘survival’ (Therneau 2021) and ‘survminer’ (Kassambara et al. 2020) to test for differences in the probability of emergence based on density, elevation, and their interaction.

To characterize behavioral differences between the field and the laboratory, we constructed linear mixed models using the ‘lmer’ function in the ‘lme4’ package (Bates et al. 2011) with frog ID as a random effect and trial (field vs laboratory), density, elevation, and their interactions as fixed effects. We used Type III ANOVAs to generate test statistics and p-values for main effects and interactions. To prevent overfitting, we first ran models with all possible two- and three-way interactions and removed interactions when non-significant. Where appropriate, we used the ‘emmeans’ package (Lenth et al. 2023) for pairwise post-hoc comparisons. We calculated Kendall’s tau correlation coefficients and test statistics between behaviors and RMR using the cor.test function from the ‘Hmisc’. We interpreted correlation coefficients following the criteria ≈0.1 as a small effect, ≈0.3 as a moderate effect, and values >0.5 as a strong effect (Garamszegi et al. 2013).

Finally, we again used linear models to test whether correlations between behaviors, and between RMR and behavior, depended on density and elevation at collection sites. We ran the latter tests using measures of behavior in the laboratory as these behavioral assays occurred closer in time and under the same conditions as RMR measurements.

## Results

### Field Behavioral Assays

For exploration, we found statistically significant main effects of density (F_1,37_= 4.91, p= 0.033), elevation (F_1,37_= 5.38, p= 0.026), and their interaction (i.e., site type; F_1,37_= 6.26, p= 0.017) (Table 1, Figure 1a). While the effect sizes of density (η^2^<0.01) and elevation (η^2^=0.01) were small, the effect of the interaction was large (η^2^=0.14). For activity, we found a trend toward statistical significance of density (F_1,36_= 3.34, p= 0.072) and the interaction between density and elevation (F_1,36_= 3.38, p= 0.074) (Table 1, Figure 1b). The effect sizes of density (η^2^<0.01) and elevation (η^2^<0.01) were small, but the effect of the interaction was moderate (η^2^=0.09). For emergence, there was also a trend toward significance for the effect of elevation (X^2^= 3.29, p= 0.070), with low elevation frogs emerging more quickly (Table 1, Figure 1c). However, pairwise comparisons were not significant in Tukey corrected, post hoc comparisons for any behavior (exploration, activity, or emergence).

**Figure 1.**
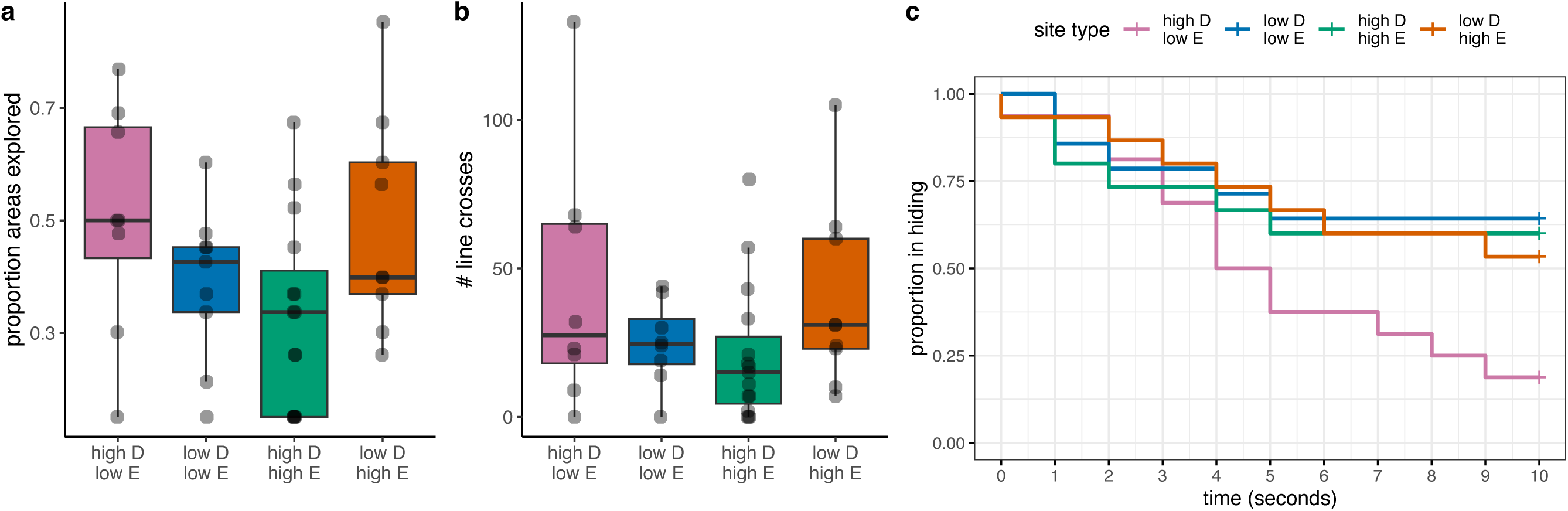
Behavior across high- and low-density (D) and elevational (E) populations in the field. Key findings include (a) a significant interaction of density and elevation on exploration, (b) a trend toward statistical significance on activity, and (c) no statistically significant differences on probability to emerge (boldness). Complete statistical results are in Table 1.

**Table 1.** Model outputs for behavioral and resting metabolic rate (RMR) differences in the field and the laboratory.

|  | FIELD |  |  |  |  |  | LAB |  |  |  |  |  |  |  |
| --- | --- | --- | --- | --- | --- | --- | --- | --- | --- | --- | --- | --- | --- | --- |
|  | <u>exploration</u> |  | <u>activity</u> |  | <u>emergence</u> |  | <u>exploration</u> |  | <u>activity</u> |  | <u>emergence</u> |  | <u>RMR</u> |  |
|  | F<sub>1,37</sub> | p | F<sub>1,36</sub> | p | X<sup>2</sup> | p | F<sub>1,52</sub> | p | F<sub>1,52</sub> | p | X<sup>2</sup> | p | F<sub>1,50</sub> | p |
| **density** | 4.91 | 0.033 | 3.43 | 0.072 | 0.02 | 0.891 | 0.00 | 0.989 | 0.08 | 0.772 | 0.06 | 0.812 | 0.74 | 0.393 |
| **elevation** | 5.38 | 0.026 | 2.83 | 0.101 | 3.29 | 0.070 | 1.51 | 0.225 | 0.57 | 0.453 | 3.08 | 0.079 | 0.32 | 0.571 |
| **density\*elevation** | 6.26 | 0.017 | 3.38 | 0.074 | 2.19 | 0.140 | 0.03 | 0.874 | 0.07 | 0.794 | 1.28 | 0.259 | 3.01 | 0.090 |

### Laboratory Behavioral Assays

After acclimation to the laboratory, we found no differences in exploration or activity based on density, elevation, or their interaction (Table 1, Figure 2a,b). The effect of elevation on emergence time had a trend toward significance in the lab as in the field (X^2^= 3.08, p= 0.079) (Table 1), though we note very few frogs emerged at all (5/54 frogs). Effect sizes were small to negligible for all models.

**Figure 2.**
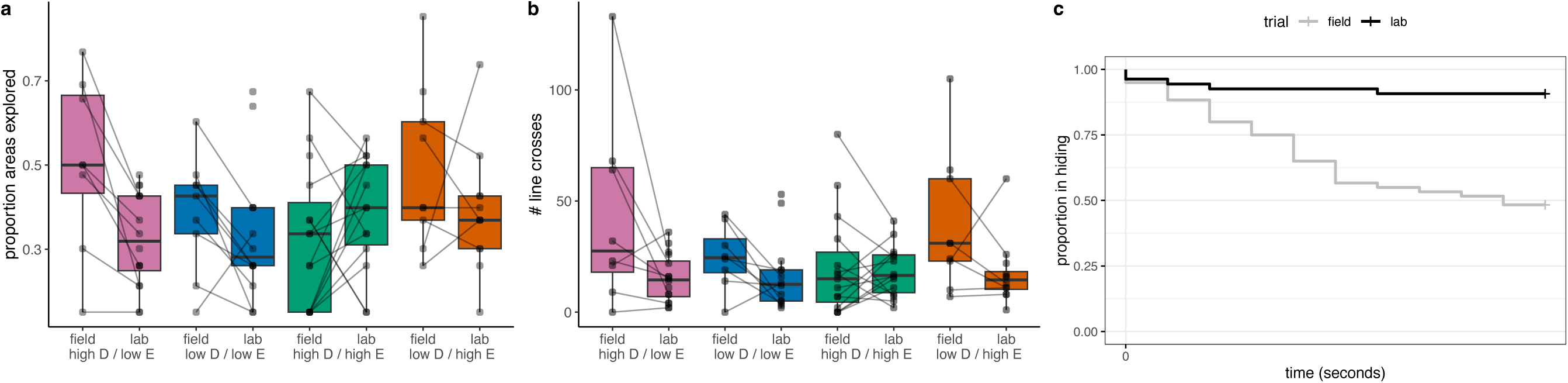
Behavioral changes from the field to the laboratory. We observed no behavioral differences based on population of origin density or elevation after acclimation to the laboratory. However, we found overall decreased (a) exploration and (b) activity, and (c) a lower propensity to emerge in the laboratory as compared to the field. For exploration and activity, this shift was particularly pronounced in high density, low elevation populations. Lines in (a) and (b) connect field and laboratory observations for the same individuals.

### Comparison of Field and Laboratory Behaviors

Comparing behavior for the same individuals in field and laboratory assays, we observed significant main effects of trial (field vs lab) on exploration (F_1,43_= 5.41, p= 0.025) and activity (F_1,47_= 7.54, p= 0.008) (Figure 2b; Table 2). No two-way interactions were significant, but the three-way interaction between trial, density, and elevation was significant for exploration (F_1,43_= 5.12, p= 0.029) and approaching significance for activity (F_1,43_= 3.80, p= 0.057), and the effect sizes were moderate (exploration: η^2^=0.11; activity: η^2^=0.07). These interactions were driven by the fact that behavioral shifts from the field to the lab were greater for some site types than others (Table 3, Figure 2a,b). Specifically, frogs from high-density/low-elevation sites showed the largest shift in behavior (exploration: t_51_= 2.79, p= 0.0046; activity: t_46_= 2.46, p= 0.018).

**Table 2.** Model outputs for comparison of behavior in the field versus the laboratory.

|  | <u>exploration</u> |  | <u>activity</u> |  | <u>emergence</u> |  |
| --- | --- | --- | --- | --- | --- | --- |
|  | F | p | F | p | X<sup>2</sup> | p |
| **trial** | 5.41 | 0.023 | 7.54 | 0.008 | 3.01 | 0.083 |
| **density** | 0.22 | 0.640 | 0.27 | 0.061 | 0.00 | 0.941 |
| **elevation** | 0.02 | 0.890 | 0.00 | 0.954 | 2.17 | 0.141 |
| **trial\*density** | 0.06 | 0.813 | 0.50 | 0.483 | 0.06 | 0.813 |
| **trial\*elevation** | 3.52 | 0.067 | 1.26 | 0.268 | 0.00 | 0.999 |
| **density\*elevation** | 3.46 | 0.069 | 2.15 | 0.148 | 1.50 | 0.220 |
| **trial\*density\*elevation** | 5.11 | 0.029 | 3.80 | 0.057 | 0.00 | 0.999 |

**Table 3.** Post hoc comparison of within site type behavioral differences in the lab versus field.

|  | FIELD vs LAB |  |  |  |
| --- | --- | --- | --- | --- |
|  | <u>exploration</u> |  | <u>activity</u> |  |
|  | t | p | t | p |
| high density / low elevation | 2.97 | 0.005 | 2.46 | 0.016 |
| high density / high elevation | -1.18 | 0.245 | -0.61 | 0.549 |
| low density / low elevation | 1.07 | 0.291 | 1.25 | 0.219 |
| low density / high elevation | 1.36 | 0.178 | 1.99 | 0.051 |

The effect of trial on emergence probability showed a non-significant trend in the full model (X^2^= 3.01, p= 0.083) (Table 2) and a significant effect (X^2^= 18.48, p<0.0001) when all non-significant two-and three-way interactions were removed (Figure 2c). In general, frogs explored less, moved less, and were less likely to emerge in the laboratory as compared to the field (Figure 2).

Activity and exploration were strongly correlated in both the field (tau= 0.82, z= 7.25, p<0.001) and the lab (tau= 0.71, z= 7.38, p<0.001), with the strength of this relationship was significantly different based on trial context (F_1,92_= 4.9, p= 0.029) (Figure 3). Neither exploration nor activity was correlated with time to emerge in the field (exploration: tau= -0.06; activity: tau= 0.004) or the lab (exploration: tau= 0.04; activity: tau= 0.03).

**Figure 3.**
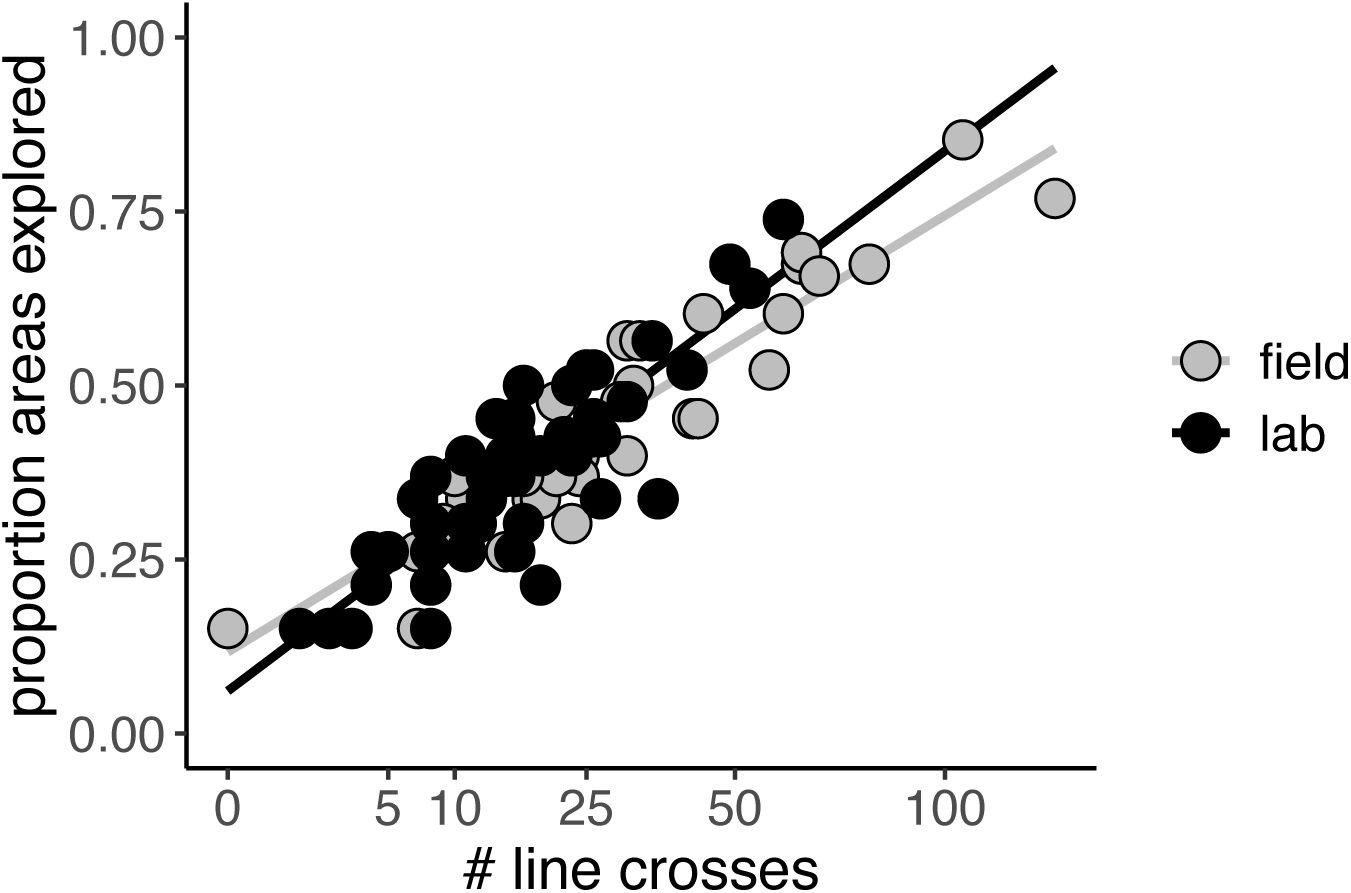
Activity and exploration are strongly correlated. While we expected a positive correlation between activity and exploration due to assay design, these metrics are not perfectly correlated, and we tested for variation between populations or origin and between field and laboratory trials. While we found strong, significant correlations under both conditions, the strength of the relationship was significantly different in the field (tau=0.82; grey line) and the laboratory (tau=0.71; black line). Activity (# line crosses; x-axis) is shown on a square root adjusted scale to improve separation and visualization of points at the low end.

Due to known effects of temperature and humidity on behavior, and variation in these metrics between the lab and the field, we tested for effects of temperature and humidity on behavior within and across contexts. We found no significant effects of either temperature or humidity on any behavior.

### Physiological Assays

We found no main effect of density (F_1,50_= 0.74, p= 0.393) or elevation (F_1,50_= 0.32, p= 0.571) on RMR after acclimation to the laboratory, but a non-significant trend in their interaction (F_1,50_= 3.00, p= 0.089) (Figure 4a). The effect sizes of density (η^2^=0.01) and elevation (η^2^<0.01) were small, but the effect of the interaction was moderate (η^2^=0.06). However, pairwise differences between site types were not significant in Tukey corrected, post hoc comparisons. While there were no substantial differences in RMR based on population of origin, RMR moderately predicted exploration (tau= 0.27, F_1,50_= 12.03, p= 0.001) (Figure 4b) and activity (tau= 0.25, F_1,50_= 7.53, p= 0.008) (Figure 4c), but not propensity to emerge (tau= -0.02, F_1,50_= 0.20, p= 0.66).

**Figure 4.**
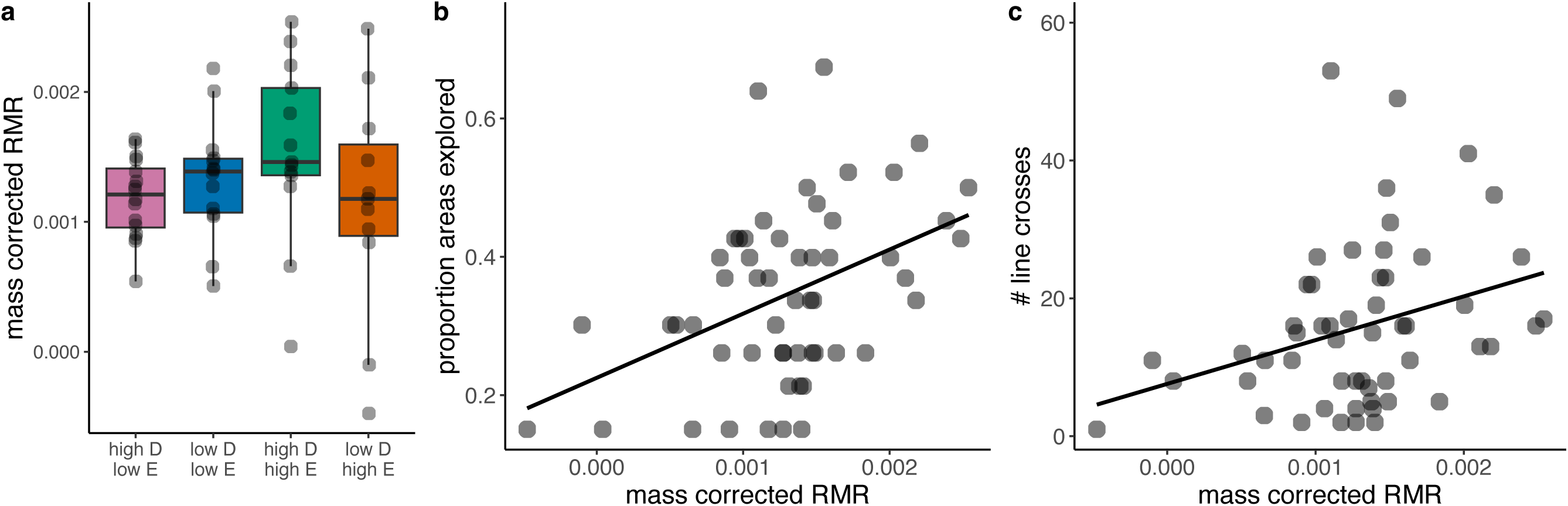
Resting metabolic rate (RMR) and behavior. (a) The interaction between population of origin density and elevation on RMR trended toward statistical significance (Table 1) but all post hoc pairwise comparisons between site types were non-significant. There were moderate, significant correlations between RMR and (b) exploration (tau=0.27) and (c) activity (tau=0.25) (behavioral data from laboratory trials only). RMR is represented as the volume of CO_2_ produced per minute per gram of wet mass (uL CO_2_/min/g).

## Discussion

During their expansion across Hawaii, coquí have experienced changes in ecological conditions, notably extreme increases in population densities and climatic fluctuations associated with colonizing higher elevations. Previous research has characterized the consequences of these shifting conditions on ecosystem dynamics, morphology, and physiology (Beard et al. 2008; Rollins-Smith et al. 2015; Haggerty 2016; O’Neill et al. 2018; Marchetti et al. 2023; Marchetti et al. 2024), but behavior remains relatively understudied in this context (but see Marchetti and Beard 2021). To address this gap, we conducted a series of experiments, examining variations in behavior across environmental and experimental contexts. We found that behavior varied based on interactions between density and elevation across the invasive range. However, these field-based differences disappeared after acclimation to the laboratory, suggesting behavioral variation in the field is primarily driven by behavioral plasticity rather than local adaptation. Additionally, we found that physiology predicted behavior in the laboratory. Below, we discuss the implications and limitations of our findings.

Our initial prediction was that behavior would vary with density if certain individuals were more likely to expand outward from high-density population centers toward low-density population edges, and that this effect might be modulated by the challenges associated with moving up in elevation. In the field, we found a trend toward faster emergence time at high densities, in line with the prediction that individuals living at high densities are bolder, perhaps conferring an advantage in acquiring food and mates in a competitive environment. For exploration and activity, we found a significant interaction between density and elevation on exploration, and a trend toward the interaction of density and elevation influencing activity.

Importantly, effect sizes indicated that neither variable alone predominated, but that the effects of density differ across elevations. For example, interactions could arise if density dependent differences associated with competitive abilities are modulated by the physiological constraints imposed by higher elevations. We note that our sample sizes for exploration and activity were relatively small. Nonetheless, statistical analyses indicated sufficient power to detect medium to large effects, suggesting that—if additional behavioral differences between populations do exist—they are subtle. Furthermore, previous studies in amphibians have primarily compared invasive and native populations, where behavioral differences might be more pronounced. We suggest this is an interesting avenue for future comparisons in coqui. In sum, while the subtle effects we identify provide only weak evidence for behavioral differences associated with invasive range expansion in Hawaiian coqui, they underscore the potentially complex interactions of multiple factors influencing behavior.

Although we found little evidence for population differences in the field, by comparing behavior of the same individuals in the field and after acclimation to the laboratory, we found strong evidence of behavioral changes in a novel environment. Following acclimation to the laboratory, frogs from all populations exhibited reduced activity, exploration, and boldness. We propose two potential explanations for these behavioral shifts. First, decreased activity and exploration may signal a shift towards cryptic behavior, which is a key factor in the success of invasive species, enabling them to evade predation, reduce competition, and thrive in new environments (Jarić et al. 2019). Alternatively, this behavioral shift may result from a reduced need or motivation to move and explore under laboratory conditions where there is an absence of resource competition and mating opportunities. Although we cannot distinguish these alternatives, uniform behavior in a novel environment indicates a generalized strategy across individuals from different site types, suggesting environmentally induced behavioral responses—rather than local adaptation—drive behavioral variation among Hawaiian coquí.

Behavioral plasticity may enhance invasion success by enabling rapid responses to new environmental conditions, even in the absence of genetic variation (Wright et al. 2010). Indeed, another study in Hawaiian coquí found that behavioral plasticity in microhabitat use may mediate colonization of higher elevation sites (Marchetti et al. 2024), providing an explanation for a lack of physiological changes among coquí from different elevations in some studies (Haggerty 2016; Marchetti et al. 2024), and underscoring the potential for a combination of genetic and environmental influences underlying traits across coquí’s native and invasive range.

In addition to single behaviors, we were interested in correlations among our behavioral metrics. Chapple et al. (2012) suggest that collections of correlated behaviors could enhance invasive success and be classified as an ’invasion syndrome’ (Chapple et al. 2012). For example, correlations between aggression and activity in invasive crayfish (*Pacifastasus leniusculus*) at higher densities facilitate their dominance over native species in the communities they invade (Pintor et al. 2009). We found that exploration and activity were strongly correlated in both the lab and the field, although there was a significant decrease in the strength of this relationship in the laboratory. We expected an overall positive relationship between exploration and activity, but note that variation in the magnitude of this relationship is possible, as evidenced by our findings. Though additional work is needed, the strength of this relationship could reflect the commonly recovered activity:exploration behavioral syndrome (for a review in frogs see Kelleher et al. 2018). Neither exploration nor activity were correlated with time to emerge, our measure of boldness. However, because emergence probability was overall low, especially in the laboratory, it is difficult to draw conclusions. Additionally, commonly used measures of boldness are not always correlated with one another (Carter et al. 2013; Yuen et al. 2017), and here too, additional work is needed. In sum, our findings here provide starting points for future work characterizing additional behaviors and the same behaviors in alternative assays.

In addition to correlations among behaviors, correlations between behavior and physiology also play a role in invasive success (Candler and Bernal 2015; Damas-Moreira et al. 2019). Coquí in Puerto Rico and Hawaii show physiological variation across elevational gradients (Chaparro 2023; Marchetti et al. 2023; Marchetti et al. 2024) and in response to manipulations in the lab (Haggerty 2016; Marchetti et al. 2023). Therefore, we were interested in whether and how such physiological differences might be associated with behavioral variation.

We measured resting metabolic rate (RMR) after acclimation to the laboratory to ask whether we could still detect differences based on population of origin and whether RMR was associated with behavior. We found a trend toward an interaction of density and collection site elevation on RMR following acclimation to laboratory conditions. Notably, these effects mirrored behavioral findings, where the effects of density were modulated by elevation. As we did not measure RMR in the field, we cannot say whether RMR differed among site types in the field. However, we also found a positive association between RMR and exploration and activity in the laboratory.

Taken together with those of others, our findings here demonstrate associations between metabolic rate and behavior, and suggest that both change plastically in response to environmental conditions across the invasive range of coqui in Hawaii.

### Conclusions

Our findings highlight the relationships between behavioral traits, physiological factors, and environmentally induced changes in an impressive invader. Collectively, our findings underscore the ability of coquí to thrive in diverse ecological conditions and suggest that behavioral and physiological plasticity play a role in their success as invaders. This study contributes to the growing body of research on the role of behavior in invasive potential.

## Acknowledgements

We thank the members of the Fischer Laboratory for their help with frog care, input on data analysis, and feedback on previous versions of the manuscript. For their assistance with equipment and technical support, we thank Dr. Jen Moss and Reed Crawford, as well as the Illinois Natural History Survey and the University of Illinois Urbana Champaign (UIUC) College of Agricultural, Consumer, and Environmental Sciences for providing technical equipment funding. We thank Michael R. Britton for advice on metabolic rate data collection and analysis. We thank Jack Marchetti, Drs. Karen H. Beard and Susannah S. French, and residents and employees of the Hawaii Division of Wildlife for their assistance with navigating the permitting process and identifying collection sites. Our work would not be possible without the support of the Division of Animal Resources at UIUC.

## Data Availability

Data analyzed in the current manuscript are included as supplemental materials.

## Author Contributions

Conceptualization: KMS and EKF. Methodology: KMS, DE, ALC, and FOH. Formal analysis: KMS and EKF. Investigation: KMS, DE, ALC, and FOH. Data curation: KMS and EKF. Writing – Original draft: KMS and EKF. Writing – Review & Editing: KMS, DE, ALC, FOH, and EKF. Visualization: KMS and EKF. Project administration: EKF. Funding acquisition: KMS and EKF.

